# Chances and challenges of machine learning based disease classification in genetic association studies illustrated on age-related macular degeneration

**DOI:** 10.1101/867697

**Authors:** Felix Günther, Caroline Brandl, Thomas W. Winkler, Veronika Wanner, Klaus Stark, Helmut Küchenhoff, Iris M. Heid

## Abstract

Imaging technology and machine learning algorithms for disease classification set the stage for high-throughput phenotyping and promising new avenues for genome-wide association studies (GWAS). Despite emerging algorithms, there has been no successful application in GWAS so far. We established machine learning based disease classification in genetic association analysis as a misclassification problem. To evaluate chances and challenges, we performed a GWAS based on automated classification of age-related macular degeneration (AMD) in UK Biobank (images from 135,500 eyes; 68,400 persons). We quantified misclassification of automatically derived AMD in internal validation data (images from 4,001 eyes; 2,013 persons) and developed a maximum likelihood approach (MLA) to account for it when estimating genetic association. We demonstrate that our MLA guards against bias and artefacts in simulation studies. By combining a GWAS on automatically derived AMD classification and our MLA in UK Biobank data, we were able to dissect true association (*ARMS2*/*HTRA1, CFH*) from artefacts (near *HERC2*) and to identify eye color as relevant source of misclassification. On this example of AMD, we are able to provide a proof-of-concept that a GWAS using machine learning derived disease classification yields relevant results and that misclassification needs to be considered in the analysis. These findings generalize to other phenotypes and also emphasize the utility of genetic data for understanding misclassification structure of machine learning algorithms.

## INTRODUCTION

Imaging technology allows for non-invasive access to detailed disease features in large studies and genome-wide association studies (GWAS) on such disease phenotypes can be expected to accelerate knowledge gain. However, image-based disease classification can be challenging for large sample sizes due to time-intensive, tiresome manual inspection. This limitation can be overcome by automated disease classification via machine learning and particularly deep learning algorithms. Such emerging approaches^1^ can classify diseases effortlessly also for huge sample sizes as needed for GWAS or other -omics approaches.

Deep learning algorithms require enormous input data with available gold standard classification, in order to “learn” classification reliably. Once trained and tested, the algorithms can be applied to external image data, but they cannot critically reflect unusual findings or incorporate unforeseen aspects, for which the human eye and brain has un-met capability. At the current time, the input data to train algorithms is limited and often specific to a certain setting (e.g. patients from a clinic). Some characteristics that appear useful for disease classification in one setting might be misinterpreted in another, which can hamper transferability of trained models; a topic discussed as dataset shift or domain shift^2–4^. Most predictions of deep learning algorithms for image-based disease classification will be error-prone and the structure of misclassification will generally be unknown. When using automated disease classification as outcome for association analyses and GWAS, the underlying response misclassification is usually unaccounted for, giving rise to biased effect estimates and potentially false-positive associations^5–7^. Extent and structure of the misclassification process can be assessed by *internal validation data*, i.e. a subset of participants with both automated and gold standard classification, which can also be utilized to account for response misclassification in statistical models^7,8^.

At present, it is unclear whether machine learning based disease classification is of any utility for association analyses, particularly for detecting disease signals in GWAS. We thus set out to evaluate machine learning derived disease classification in GWAS on the example of age-related macular degeneration (AMD) and we developed a statistical approach accounting for the implied response misclassification. AMD is an ideal role model, as a common disease ascertained via imaging of the central retina^9^ and with particularly strong known genetic effects^10^. The manual grading of images for AMD requires a substantial effort by trained staff and is currently an obstacle for homogeneous disease classification within and across large studies. For example, in UK Biobank^11^, >135,000 color fundus images are available for >68,000 study participants, but there is no manually classified AMD available so far. Several machine learning algorithms have been emerging to classify AMD: some show promising performance, but still yield misclassified predictions, have acknowledged issues due to domain shift or insufficient sample size for training, or they lack validation in external studies^12–15^. So far, there is no GWAS on fundus image ascertained AMD available in UK Biobank, manually classified or machine learning based.

## MATERIALS AND METHODS

### Machine learning based disease classification in GWAS as misclassification problem

We consider a binary disease Y, for which each individual has a true status of disease (disease yes/no). A *gold standard* classification often involves manual grading of medical images via trained medical staff, which is considered here to correspond to the true disease classification. When applying a trained machine learning algorithm on medical images, we yield an automated disease classification Y* for each individual. For an individual i with true disease status Y_i_ = y_i_, the classification 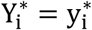 can either be correct or error-prone.(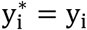, or 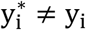). If a gold standard classification is available (for at least a subset of study participants, internal validation data), the performance of the algorithm can be quantified by cross-tabulation of the observed error-prone y* and the gold-standard classification y across all participants in the validation sub-study (confusion matrix); the (mis-)classification process can be characterized by classification probabilities P(Y* = k|Y = l), for l, k ∈ {0,1}. For l = k = 1 and l = k = 0, these probabilities correspond to the sensitivity and specificity of the algorithm, respectively.

In the following, we focus on *bilateral diseases* due to our motivating example of an eye disease (AMD): for each individual i, two entity-specific binary disease variables Z_1i_, Z_2i_ ∈ {0,1} (here: AMD per eye) are used to define the binary person-specific disease status as the “worse-entity disease status” Y_i_ ≔ max(Z_1i_, Z_2i_), corresponding to “AMD in at least one eye” versus “AMD in none of the two eyes” in our example. The error-prone machine learning based classification of entity-specific disease 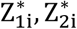, will propagate to an error-prone person-specific disease status, 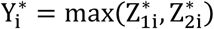, when compared to the manually graded; “true” Y_i_.

We were interested in evaluating the potential and consequences of such automatically classified disease in GWAS. The standard approach in GWAS is logistic regression for modelling the association of a genetic variant (observed as genotypes ∈ {0,1,2} or imputed allelic dosages ∈ [0,2]) with a binary disease status, usually adjusted for other covariates like age, sex, and genetic principal components; Wald-tests are used to test for genetic association, accounting for multiple testing by judging at a Bonferroni-corrected significance level of p<5 × 10^−8^. When the association of the genetic variant with the true disease status Y (here: manually classified persons-specific AMD) follows a logistic regression model, the usage of the error-prone disease status Y* (here: automatically derived person-specific AMD) in the logistic regression will lead to a mis-specified model (*naïve association analysis*) with known consequences of decreased power, biased genetic association estimates, and potentially false-positive associations^5–7^.

### MLA to adjust for response misclassification in bilateral disease

While there are methods available to account for response misclassification for classic diseases in standard logistic regression^5–7^, there is currently no methodology readily available for bilateral disease. As described previously^16^, the conceptual challenge here is to account for two types of misclassification: (i) the entity-specific misclassification that propagates to an error-prone person-specific disease status, where the person-specific disease status is used in the association analysis, and (ii) a person-specific misclassification from a missing disease status in one of the two entities. We thus developed an MLA to account for the fact that we are using an error-prone response 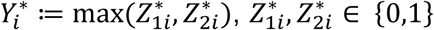, in the association analysis, while the true disease *Y*_*i*_ ≔ max(*Z*_1*i*_, *Z*_2*i*_), *Z*_1*i*_, *Z*_2*i*_ ∈ {0,1}, is assumed to follow a logistic regression model.

Details are provided in **Appendix A**. The general idea of the MLA is to factorize the likelihood of the observed, error-prone response data into two parts, the model for the association between risk factor and true (but in general unobserved) response (*true association model*) and a model for the misclassification process (*misclassification model*). We adapted this well-established methodology for analyzing misclassified binary response data^7,8^ to the scenario of bilateral disease with a “worse-entity” disease definition (i.e. the person-specific disease status is defined as the status of the worse entity). Under the assumption of independent misclassification for the observed disease in the two entities 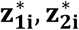 of an individual i, we derive

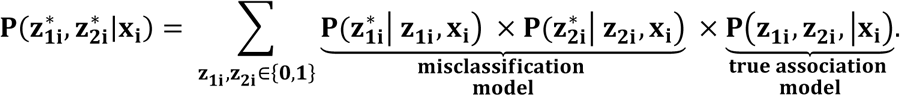

The *misclassification model* is characterized by the sensitivity and specificity of the entity-specific classification process; the *true association model* is the assumed logistic regression model for the person-specific disease status. When internal validation data is available, the parameters of both models can be estimated jointly by optimizing a likelihood with different contributions of participants with only the error-prone response and participants in the validation data with true and error-prone response available.

Our developed approach allows us to adjust for both the entity-specific misclassification from an automated classification and the misclassification of the person-specific status when one entity is ungradable. Altogether, we model four parameters in the MLA: (i) the conditional probability of worse-entity disease given the covariate of interest, (ii) the probability of disease in both entities conditional on the disease in at least one entity (to adjust for missing information of one of two entities), as well as (iii) the sensitivity and (iv) the specificity of the entity-specific misclassification process. For each parameter, the conditional probabilities are modeled using the logistic function (as in standard logistic regression) allowing for a dependency on a parameter-specific set of person-specific covariates. An open source R implementation is available (**Web Resources**).

### Simulation study to investigate the performance of the MLA

We repeatedly simulated association data for a standard normal covariate X and a (true and error-prone) binary outcome of a *bilateral* disease. To do this, we (1) sampled the true, person-specific worse-entity status associated with X, (2) derived the true entity-specific disease status (e.g. manual eye-specific AMD classification) given assumptions, (3) sampled the entity-specific error-prone disease status (e.g. automated AMD classification), and (4) derived an error-prone, person-specific disease status. Afterwards, we removed the true disease status for most individuals, yielding only a subset with both true and error-prone disease status available (validation data). In different simulation scenarios, we varied sensitivity and specificity of the entity-specific classification. Classification probabilities were either constant for all individuals (non-differential misclassification) or varying with X (differential misclassification). We also varied the fraction of individuals with missing classification in one of two entities. Data was sampled with or without an effect of X on the true person-specific response Y (β_Y_ϵ{0,1}, log OR) and on the probability δ of having disease in both entities given disease in at least one entity (β_δ_ϵ{0,1}, log OR). We estimated the covariate effect using the naive analysis (logistic regression, which ignores misclassification) and the developed MLA1 and MLA2 accounting for response misclassification without (MLA1) and with allowing (MLA2) for differential misclassification, respectively. To compare the performance of the naïve analysis and the derived MLA, we investigated the distribution of effect estimates 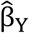 across simulation runs, computed the mean squared error of estimates relative to true effects, frequencies of rejected tests for no association, and coverage frequencies of 95%-confidence intervals. A detailed description of the simulation study, data sampling, and estimated models is given in **APPENDIX B**.

### UK Biobank study information and data

UK Biobank recruited ∼500,000 individuals aged 40-69 years from across the United Kingdom. Genetic data is available from the Affymetrix UK Biobank Axiom Array imputed to the Haplotype Reference Consortium^17^ and the UK10K haplotype resource^18^ (details described elsewhere^11^). The UK Biobank baseline data contains 135,500 fundus images of 68,400 individuals. The images are taken with the Topcon 3D OCT-1000 Mark II sytem with a field angle of 45° without application of mydriasis^19^. The images can be utilized for automated or manual AMD classification, however, there is no image-based AMD classification publicly available so far.

### AMD classification in UK Biobank derived from a machine learning algorithm and manually

We performed an automated AMD classification for 68,400 individuals with available fundus images in UK Biobank with additional manual classification in a subset of 2,013 participants as described in the following.

In epidemiological studies, AMD is usually classified per eye via manual grading of color fundus images by trained graders using established classification systems. One such system is the Age-Related Eye Disease Study (AREDS) 9-step Severity Scale^20^, which defines early AMD combining a 6-step drusen area scale with a 5-step pigmentary abnormality scale and is therefore particularly detailed and time-consuming when applied manually. Another more recent system is the Three Continent AMD Consortium Severity Scale (3CC)^9^, which defines early AMD based on drusen size, drusen area and presence of pigmentary abnormalities and is thus more practical to apply manually. While the definition of “advanced AMD” is fairly robust across systems, each system defines “early” or “intermediate” AMD differently, but provides a clear assignment strategy to “no”, “early/intermediate” or “advanced AMD” (or “no” and “any AMD”).

To obtain an eye-specific AMD status for the 135,500 images of the UK Biobank (≤ 1 image per eye; 67,100 individuals with images for both eyes, 1,300 with image for only one eye), we applied a published convolutional neural network ensemble^14^ to the fundus images following recommendations of the authors (**Web Resources**). The ensemble was trained to classify each image into the AREDS 9-step severity scale or three additional categories for advanced AMD (GA, NV, mixed GA+NV, “AREDS 9+3 steps”) or “ungradable”. From this, we derived the person-specific automated AMD status as the AMD status of the worse eye (i.e. the higher score of the ARED9+3) or as the status of the only eye, if applicable. We collapsed AREDS AMD severity steps 2-9 or any of the 3 advanced AMD categories to “any AMD”.

To generate internal validation data, we selected a subset of UK Biobank individuals for additional manual grading. When randomly sampling participants, one would expect to catch only a few AMD individuals; we thus enriched the validation sample with persons likely to be affected by AMD or likely to be unaffected: (i) persons with high genetic risk score for AMD based on the known 52 variants for advanced AMD^10^ (> 99^th^ percentile, n=829), (ii) persons with low genetic risk score (<1^st^ Percentile, n=828), and (iii) persons with self-reported AMD not already selected (n=356). The machine learning based AMD classification was not used to select individuals into the validation subset. The selected 2,013 individuals were manually classified for AMD according to the 3CC^9^ system by a trained ophthalmologist (five AMD categories, 1 for no AMD, 3 for early, 1 for advanced AMD, and 1 “ungradable”). We collapsed the five AMD categories to “any AMD”, “no AMD”, or “ungradable” and derived eye-specific as well as person-specific confusion matrices based on the detailed (AREDS 9+3 and 5-category 3CC) and collapsed classifications. To conduct the GWAS with automatically derived “any AMD”, we restricted the data with available automated AMD classification to unrelated individuals of European ancestry with valid GWAS data (see below), and derived the confusion matrices also for the restricted validation data.

### Genetic association analyses for AMD without and with accounting for misclassification

We performed a GWAS on the automatically derived “any AMD” versus “no AMD” in unrelated UK Biobank participants (relatedness status > 3^rd^ degree) of European ancestry (self-report “White”, “British”, “Irish” or “Any other white background”) as recommended^21^. For each variant, we applied a standard logistic regression model (i.e. the naïve analysis ignoring misclassification in the automatically derived AMD status) under the additive genotype model and applied a Wald-test as implemented in QUICKTEST^22^. We included age and the first two genetic principal components as covariates. We excluded variants with low minor allele count (MAC<400, calculated as MAC = 2 × N × MAF, sample size N, minor allele frequency MAF) or with low imputation quality (rsq<0.4) yielding 11,567,158 analyzed variants. To correct for potential population stratification, we applied a Genomic Control correction (lambda = 1.01 based on the analyzed variants excluding the 34 known AMD loci)^23^.

We selected genome-wide significant variants (P_GC_<5.0×10^−8^), clumped them into independent regions (≥500kB between independent regions) and selected the variant with lowest P-value in each region (“lead variant”). We also selected the 21 of the 34 reported lead variants from the established advanced AMD loci, for which we had ≥80% power to detect them in a UK Biobank sample size of 3,544 cases and 44,521 controls with nominally significance - under the assumption that the reported effect sizes for advanced AMD were the true effect sizes and ignoring any misclassification in the AMD classification (**APPENDIX C**). Information on linkage disequilibrium in Europeans was obtained from LDLink^24^. Enrichment of directionally consistent or enrichment of nominally significant association for the 21 reported lead variants (when compared to the reported direction literature) was tested based on the Exact Binomial test for H_0_: Prob = 0.5 or H_0_: Prob = 0.05, respectively.

To evaluate the robustness of the genetic association upon accounting for the misclassification, we applied the derived MLAs for the selected variants. For this, we modelled the conditional probability of AMD depending on age, genetic variant and two genetic principal components (as in the naïve analysis). The MLAs accounted for the misclassification of the eye-specific automated classification and for the person-specific misclassification from missing AMD status in one of two eyes. For the misclassification process of the eye-specific automated classification (quantified by sensitivity and specificity), we allowed for a linear association with age and modelled two scenarios for the association with the genetic variant: (i) no dependency (non-differential, MLA1) or (ii) linear dependency (differential misclassification, MLA2). We compared association estimates of the naive analysis with MLA1- and MLA2-analysis and judged significance at Bonferroni-corrected significance levels for a family-wise error rate of 0.05. To allow for comparisons across different models, we did not apply Genomic control correction for these comparative analyses. Additionally, we evaluated robustness of findings from the naïve analysis for the selected lead variants upon adjusting for 20 instead of 2 genetic principal components.

To follow-up on the *HERC2* lead variant finding (see Results), we quantified lightness of fundus images by calculating gray levels for the “RGB” fundus images (weighted sum of R, G and B values, 0.30*R+0.59*G+0.11*B, as implemented in IrfanView).

## RESULTS

### Linking misclassification theory to machine learning disease classification

We here establish the usage of machine learning derived disease classification in genetic association analyses as a response misclassification problem in logistic regression (**Methods**). We present a newly developed maximum likelihood approach (MLA) for *bilateral diseases* like AMD (**Methods**). This includes two versions: (1) assuming *non-differential misclassification* (MLA1, i.e. no dependency of misclassification probabilities on the covariate of interest, here the genetic variant) and (2) allowing for *differential misclassification* (MLA2, i.e. dependency on the covariate of interest). There are existing MLAs for considering response misclassification in logistic regression using internal validation data^7,8^: these MLAs refer to *classic diseases* where the misclassification is on the person-specific disease status. Our developed approach provides a general framework for bilateral diseases with entity-specific misclassification that propagates to person-specific disease misclassification. Our approach also allows for missing classification in one of two entities, which is a second source of bias in association analyses for bilateral diseases as reported previously^16^. We exemplify our approach on machine learning derived AMD compared to manually graded AMD. Since machine learning algorithms for AMD are trained on images with human manual AMD grading as benchmark, we assume the manual classification to be gold standard.

We evaluated the performance of our developed MLA1 and MLA2 in a simulation study. By this, we documented substantial bias and lack of type-I error control when the naïve analysis was applied, which was comparable to theory for classic (non-bilateral) diseases ^5,7^. We also showed our MLA1 and MLA2 to effectively remove bias and keep type-I error when specified correctly (**Table 1, APPENDIX D, Supplementary Table 1**).

**Table 1.** Simulation results on effect estimates and empirical type-I error in naïve and MLA-analysis. We evaluated the performance of naïve and MLA analysis of a quantitative covariate X and a binary bilateral disease Y, e.g. person-specific AMD, simulating various scenarios. For each scenario, we sampled 1000 data sets à 5000 individuals, 4000 with only error-prone eye-specific AMD classification, and 1000 with additional true AMD classification. Shown are performance measures from three models, naïve analysis, MLA1, or MLA2 assuming non-differential/differential misclassification regarding X, respectively, in various simulation scenarios. For the eight scenarios shown here, we assumed no association of X with δ, the probability of AMD in both eyes given ≥1 affected eye; results were similar when modelling an association of X with δ, see **Supplemental Table 1**. For each model and scenario, we report mean effect estimates 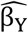, log OR per unit increase in standard-normal X, over all simulation runs, and the associated root mean squared error (RMSE), fraction of nominally significant effect estimates (% with P<0.05), and coverage frequencies of 95%-confidence intervals.

| Simulation Scenario | | | | | | $\widehat{\beta}_Y$ | | | | | | % with P<0.05 | | | Cov. Freq. | | |
| --- | --- | --- | --- | --- | --- | --- | --- | --- | --- | --- | --- | --- | --- | --- | --- | --- | --- |
| Sens | Spec | %miss. | $\beta_Y$ | $\beta_{sens}$ | $\beta_{spec}$ | Naïve | | MLA1 | | MLA2 | | Naive | MLA1 | MLA2 | Naive | MLA1 | MLA2 |
|  |  |  |  |  |  | Mean | RMSE | Mean | RMSE | Mean | RMSE |  |  |  |  |  |  |
| Non-differential misclassification |  |  |  |  |  |  |  |  |  |  |  |  |  |  |  |  |  |
| 0.9 | 0.9 | 0.25 | 0 | 0 | 0 | 0.00 | 0.03 | 0.00 | 0.04 | 0.00 | 0.04 | 5.3% | 4.6% | 4.6% | 94.7% | 95.4% | 95.4% |
| 0.9 | 0.9 | 0.25 | 1 | 0 | 0 | 0.73 | 0.27 | 1.00 | 0.05 | 1.00 | 0.05 | 100% | 100% | 100% | 0.0% | 96.5% | 96.3% |
| 0.9 | 0.9 | 0.75 | 1 | 0 | 0 | 0.69 | 0.31 | 1.00 | 0.06 | 1.00 | 0.07 | 100% | 100% | 100% | 0.0% | 94.4% | 93.5% |
| 0.8 | 0.8 | 0.25 | 1 | 0 | 0 | 0.56 | 0.44 | 1.00 | 0.06 | 1.00 | 0.07 | 100% | 100% | 100% | 0.0% | 95.0% | 95.0% |
| 0.8 | 0.9 | 0.25 | 1 | 0 | 0 | 0.68 | 0.32 | 1.00 | 0.05 | 1.00 | 0.06 | 100% | 100% | 100% | 0.0% | 97.0% | 95.9% |
| 0.9 | 0.8 | 0.25 | 1 | 0 | 0 | 0.61 | 0.39 | 1.00 | 0.06 | 1.00 | 0.06 | 100% | 100% | 100% | 0.0% | 95.3% | 94.8% |
| Differential misclassification |  |  |  |  |  |  |  |  |  |  |  |  |  |  |  |  |  |
| 0.9 | 0.9 | 0.25 | 0 | -1 | 1 | -0.38 | 0.38 | -0.46 | 0.46 | 0.00 | 0.05 | 100% | 100% | 4.7% | 0.0% | 0.0% | 95.3% |
| 0.9 | 0.9 | 0.25 | 1 | 1 | -1 | 1.14 | 0.14 | 1.39 | 0.40 | 1.00 | 0.06 | 100% | 100% | 100% | 4.8% | 0.0% | 95.1% |

Sens/Spec = average sensitivity and specificity of error-prone, eye-specific AMD classification; %miss. = fraction of randomly selected individuals with missing AMD classification in one of two eyes; $\beta_Y$ = log OR of X on true AMD, $\beta_{sens}$ = log OR of X on the sensitivity or $\beta_{spec}$ = log OR of X on the specificity of the eye-specific misclassification process, respectively.

### AMD in UK Biobank based on automated classification and validation data

We applied a published convolutional neural network ensemble^14^ to automatically derive eye- and person-specific AMD classifications for 68,400 UK Biobank participants with fundus images at baseline (135,000 eyes) (**Supplemental Table 2a**). From this, we derived eye-specific “any AMD” status (i.e. any early AMD stage or advanced AMD versus AMD-free) and person-specific “any AMD” status based on the worse eye (**Methods**). Among the 68,400 participants, 10,128 were ungradable for AMD in both eyes (i.e. missing person-specific AMD status, 14.8%), 4,870 were classified as “any AMD” and 53,402 as AMD-free (**Supplemental Table 2b)**. Among the 58,272 gradable participants (of these: 20.2% gradable only in one eye), 8.4% had AMD and 91.6% were AMD-free. This included 48,065 unrelated individuals of European ancestry with GWAS data (3,544 “any AMD” cases, 44,521 AMD-free controls; 19.8% with only one eye gradable; **Supplemental Table 2b**).

To quantify the performance of automated AMD classification, we manually classified AMD in a subset as internal validation data (4,001 images, ≤ 1 image per eye, 2,013 individuals). When comparing automated to manual (true) “any AMD” status, we found an eye-specific sensitivity of 73% and specificity of 90% in the full validation data and a person-specific sensitivity of 77% and specificity of 91% among the participants in the GWAS (**Table 2a/b**). We found no structural differences between the full validation data and when restricting to the GWAS data (1,327 individuals, **Supplemental Table 3a/b**). Both, the manual and automated classification included the category “ungradable”. Among the 4,001 eyes, 1,101 were manually ungradable, of which the automatic classification yielded 74% as ungradable as well, but classified 9% as AMD and 17% as AMD-free, which raises concerns about these classifications. In summary, we found the automated classification to yield reasonable, but error-prone results.

**Table 2.** Confusion matrices comparing manual and automated AMD classification per eye and per person. Shown are absolute numbers and conditional classification probabilities, i.e. in row i and column j, P(automated = j | manual=i) as %, with i, j=“Ungradable”, “No AMD”, “Any AMD”: **a)** for all eyes in the validation data; 4001 eyes of 2,013 individuals. **b)** For all persons in the overlap between validation data and GWAS; 1,327 persons.

| a) per eye (4,001 eyes, 2,013 individuals) |  |  |  |  |
| --- | --- | --- | --- | --- |
|  | Automated classification |  |  |  |
| Manual | Ungradable | No AMD | Any AMD | Sum |
| Ungradable | 813 (74%) | 185 (17%) | 103 (9%) | 1101 (100%) |
| No AMD | 107 (4%) | 2207 (90%) | 138 (6%) | 2452 (100%) |
| Any AMD | 20 (4%) | 103 (23%) | 325 (73%) | 448 (100%) |

| b) per person (1,327 individuals) |  |  |  |  |
| --- | --- | --- | --- | --- |
|  | Automated classification |  |  |  |
| Manual |  | No AMD | Any AMD | Sum |
| Ungradable/NA |  | 202 (79%) | 53 (21%) | 255 (100%) |
| No AMD |  | 750 (91%) | 72 (9%) | 822 (100%) |
| Any AMD |  | 58 (23%) | 192 (77%) | 250 (100%) |

NA = true AMD status based on worse eye not available, since one eye was manually ungradable and the second AMD-free

### GWAS on automated AMD classification in naïve analysis identifies two loci

While we have some idea about the extent of the misclassification from validation data and about its impact on genetic association estimates from simulations, it is unclear whether the automated any AMD classification is “good enough” for GWAS. We conducted a GWAS for person-specific automatically derived “any AMD” in UK Biobank (3,544 “any AMD” cases; 44,521 controls) applying logistic regression as usual, which is without accounting for misclassification (naïve analysis). We found 53 variants with genome-wide significance (P_GC_<5.0×10^−8^) spread across two distinct loci (defined as lead variant and proxies +/-500kB, **Figure 1a/b; Supplemental Table 4a)**: the known *ARMS2/HTRA1* locus (lead variant here rs370974631, P_GC_=3.1×10^−20^, effect allele frequency EAF=0.23) and an unknown locus for AMD near *HERC2* (lead variant rs12913832, P_GC_=4.7×10^−16^, EAF=0.23). This *ARMS2/HTRA1* lead variant was highly correlated to the reported lead variant for advanced AMD, rs3750846, and effect estimates were directionally consistent (r^2^ =0.93; **Supplemental Table 4b**). The next best known locus is the *CFH* locus, which showed close to genome-wide significance here (smallest *P-*value P_GC_=7.0×10^−7^, rs6695321, EAF=0.62): rs6695321 is in linkage disequilibrium with two reported *CFH* variants (rs61818925, rs570618: r^2^=0.63 or 0.40, D’=0.81 or 1.00, EAF=0.58 or 0.36, respectively; **Supplemental Table 4b**) suggesting that rs6695321 captures the signals of these two reported variants.

**Figure 1.**
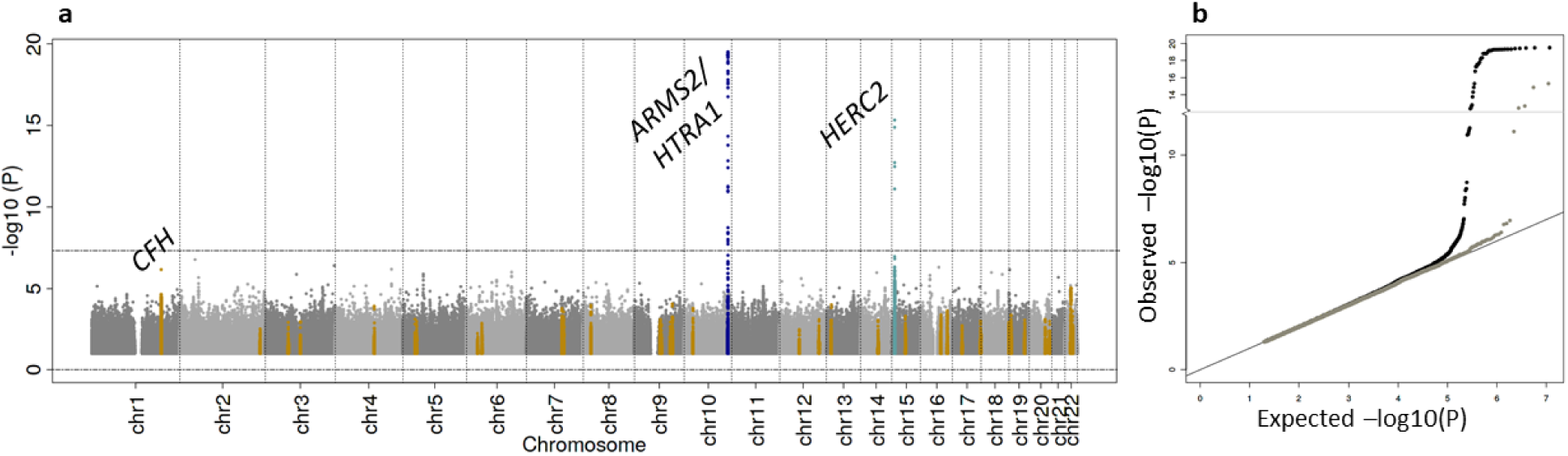
GWAS results in UK Biobank based on automatically derived “any AMD” from naïve analysis. Association analyses were conducted using the error-prone, machine learning derived AMD classification in UK Biobank participants with 3,544 “any AMD” cases and 44,521 controls via logistic regression adjusted for age and two genetic principal components, the *naïve analysis* ignoring misclassification. Shown are **a)** Manhattan Plot of 11,567,158 analyzed variants; dark blue: genome-wide significant and previously established^10^ locus, light blue: unknown genome-wide significant locus, orange: other 33 previously established loci for advanced AMD), and **b)** expected versus observed –log10 *P*-values; black: all variants, grey: all variants outside the 34 previously reported loci.

Among the reported lead variants of the 34 advanced AMD loci^10^, we had ≥80% power to detect 21 of these with nominal significance (**Supplemental Table 5**). When comparing effect sizes of these 21 variants from this analysis on “any AMD” in UK Biobank with reported effect sizes for advanced AMD, we found 15 with directional consistency (P_Bin_=0.078) and 7 with directionally consistent nominal significance (P_Bin_=4.9×10^−5^; **Figure 3a, Supplemental Table 4c**). The overall smaller effect sizes for automated “any AMD” compared to reported effect sizes for advanced AMD can be explained by a bias from misclassified automated AMD and by smaller effect sizes for early AMD merged into the definition of “any AMD”. For the other 13 of the 34 variants, we refrained from interpreting results due to lack of power in this analysis (**Supplemental Table 4c**). Results were similar when adjusting for 20 instead of 2 genetic principal components (data not shown). While the yield of only few known AMD signals in this UK Biobank GWAS may be disappointing, this is not fully unexpected given an effective sample size^25^ of 13,130 and a power estimate of ∼80% (assuming no misclassification and reported effect sizes) to detect associations with genome-wide significance for only 4 of the 34 established variants (*CFH, ARMS2/HTRA1, C3, C2/CFB/SKIV2L*, **Supplemental Table 5**).

**Figure 2.**
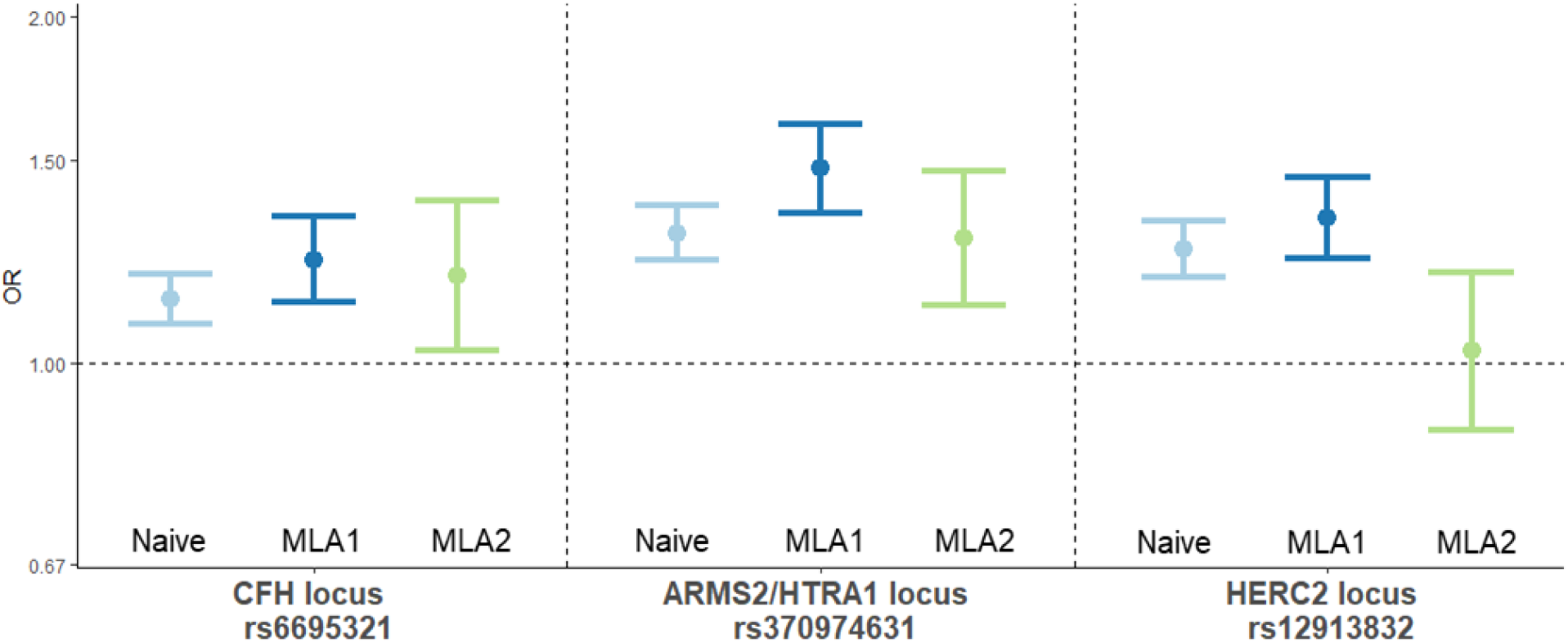
Genetic effect estimates for the 3 lead variants in UK Biobank without and with accounting for misclassification. Shown are genetic effect estimates and 95% confidence intervals for 3 lead variants from the GWAS on automated AMD classification with 3,544 “any AMD” cases and 44,521 controls from 3 models: without accounting for the misclassification; naïve analysis, light blue. With accounting for non-differential misclassification, i.e. no dependency on the genetic variant; MLA1, dark blue. And accounting for a differential misclassification, i.e. dependency on the genetic variant; MLA2, light green. Both MLAs accounted for missing AMD information in one of two eyes and a misclassification that depended on age.

**Figure 3.**
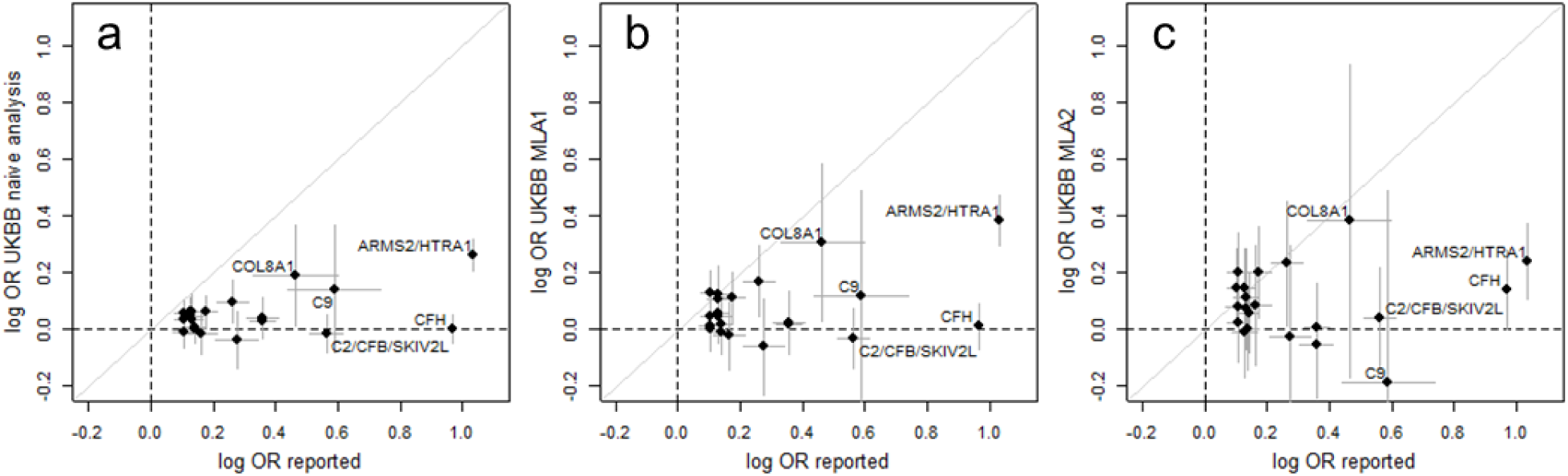
Comparison of 21 reported genetic effect estimates for advanced AMD with estimates for automatically derived “any AMD” from UK Biobank without and with accounting for misclassification. We selected the 21 reported AMD lead variants, for which we had ≥80% power to detect them in this UK Biobank sample size with nominal significance. Shown are log OR effect estimates and 95% confidence intervals reported for advanced AMD on x-axis versus UK Biobank estimates for automatically derived “any AMD” on y-axis from **a)** the naïve analysis (logistic regression ignoring misclassification, **b)** MLA1, and **c)** MLA2.

In summary, our GWAS on automated AMD in UK Biobank detected the established *ARMS2/HTRA1* locus, an unknown locus around *HERC2* with genome-wide significance, and the established *CFH* locus to some extent.

### Applying the developed MLA to account for misclassification for selected variants

Due to our simulation results and theory^5,7^, we expected our GWAS on automated (error-prone) AMD to yield biased estimates and, when the misclassification was differential towards the genetic variant, even potentially false signals. We applied our developed MLAs for 26 selected variants: (i) the 3 lead variants detected here with (near) genome-wide significance (*CFH*: rs6695321, *ARMS2/HTRA1:* rs370974631, *HERC2:* rs12913832), (ii) the 3 reported independent variants in the *CFH* locus with MAF≥5% (rs61818925, rs570618, rs10922109; 2 of these correlated to the here identified *CFH* lead variant), and (iii) the other 20 of the 34 reported lead variants^10^, for which we had reasonable power in this analysis (including 1 reported *ARMS2/HTRA1* variant correlated to here identified variant). This yielded a total of ∼23 independent variants.

Our MLAs estimated simultaneously (1) sensitivity and specificity of the eye-specific misclassification process and (2) genetic association accounting for the misclassification. With regard to sensitivity and specificity, we found (i) an overall sensitivity of 64.5% (95%-CI: 60.1%, 68.7%) and a specificity of 98.6% (98.4%, 98.8%), i.e. a false-negative “any AMD” proportion of 35.5% and a false-positive of 1.4%, (ii) no dependency of the sensitivity on any selected variant (P>0.05/(23*2)=1.09×10^−3^) and no dependency of the specificity, except for two variants: *HERC2* lead variant, rs12913832, and the reported *CFH* lead variant rs10922109 (OR_spec_=0.64, P_spec_=7.38×10^−9^ and OR_spec_=1.36, P_spec_=2.29×10^−4^, respectively; **Supplemental Table 6, Appendix E**). Therefore, we found a misclassification that was associated with some genetic variants (differential), which could induce bias into all directions and severe lack of type-I error control.

When comparing genetic association estimates from our MLA1 and MLA2 with the naïve analysis for our three detected lead variants, we found interesting patterns (**Figure 2, Supplemental Table 7a**). (i) For *CFH* and *ARMS2/HTRA1*, we found consistent effect estimates across the three analyses, with larger confidence intervals when using the more complex models MLA1 or MLA2. (ii) For *HERC2*, MLA1 yielded comparable results to the naïve analysis, but when accounting for differential misclassification (MLA2), the effect vanished (MLA2: OR=1.03, P=0.76; MLA1: OR=1.34, P=1.11×10^−12^; naïve: OR=1.26, P=4.16×10^−16^). When applying MLA1 and MLA2 to the three reported *CFH* locus variants and the further 20 of the 34 reported lead variants, we found the following (**Supplemental Table 7b/c**): (i) effect estimates for all three *CFH* variants increased when applying MLA2 compared to the naïve analysis. This was particularly interesting for the reported *CFH* lead variant rs10922109, where we now observed a nominally significant association into the reported direction (MLA2: OR=1.15, P=0.047; naïve: OR=1.00, P=0.98; **Supplemental Table 7c**). This is in line with the observed dependency of the specificity on this *CFH* variant. (ii) For the other 20 reported lead variants, many variants showed increased effect estimates by MLA2 compared to the naïve analysis (effect estimates mostly more comparable to reported effect sizes^10^; **Figure 3c**). Altogether, MLA results confirmed the *CFH* and *ARMS2/HTRA1* loci and unmasked the *HERC2* finding as false positive.

### Misclassification depended to eye and fundus image color

Interestingly, our *HERC2* lead variant, rs129138329, is precisely the variant for which the G allele was considered causal for blue eyes^26^. We were able to support this in our AugUR^27,28^ study (n=1026; reported “light eye color” for 14%, 36%, or 97% of participants with A/A, G/A, or G/G, respectively). Eye color is discussed as AMD risk factor, but the debate is on blue eyes to increase risk due to increased susceptibility to UV-radiation^29^, which is in contrast to our observation of brown eyes to increase AMD risk and a challenge for interpreting this finding. It was interesting to see the *HERC2* rs129138329 association vanish when accounting for rs129138329-associated misclassification. This was in line with the observed strong association of the specificity with this variant (OR_spec_=0.64 per A allele, **Supplemental Table 6a)** resulting in 3.0%, 1.9%, or 1.2% of false-positive AMD classifications among persons with A/A, A/G, or G/G, respectively. This notion of a larger misclassification among A/A versus G/G individuals was further supported by the larger fraction of manually ungradable images that were deemed gradable by the automatic classification among A/A versus G/G (54.5% versus 38.8%, respectively; **Figure 4**). When visually inspecting fundus images per genotype group, the images for A/A had a darker appearance than those for A/G or G/G (**Figure 4**), which we were able to quantify by means of average gray level per image of 46.4, 49.0, or 53.6, respectively. Therefore, the *HERC2* signal appeared to be an artefact due to a larger misclassification for brown eyes linked to darker fundus images. One may hypothesize that the darker eye color had reduced light exposure during fundus photography, which gave rise to darker images and more misclassified AMD-free eyes. The notion of a differential misclassification due to eye color was further supported by the fact that the full *HERC2* signal disappeared by modelling a misclassification dependency on the causal variant for eye color (rs129138329, **Supplemental Figure 1a/b**), while some signal remained when modelling a misclassification dependency on the respective *HERC2* variant in the model (**Supplemental Figure 1c**). In summary, we found the MLA2 not only to effectively remove the artefact signal of the naïve GWAS, but also to help understand the dependencies of the misclassification.

**Figure 4.**
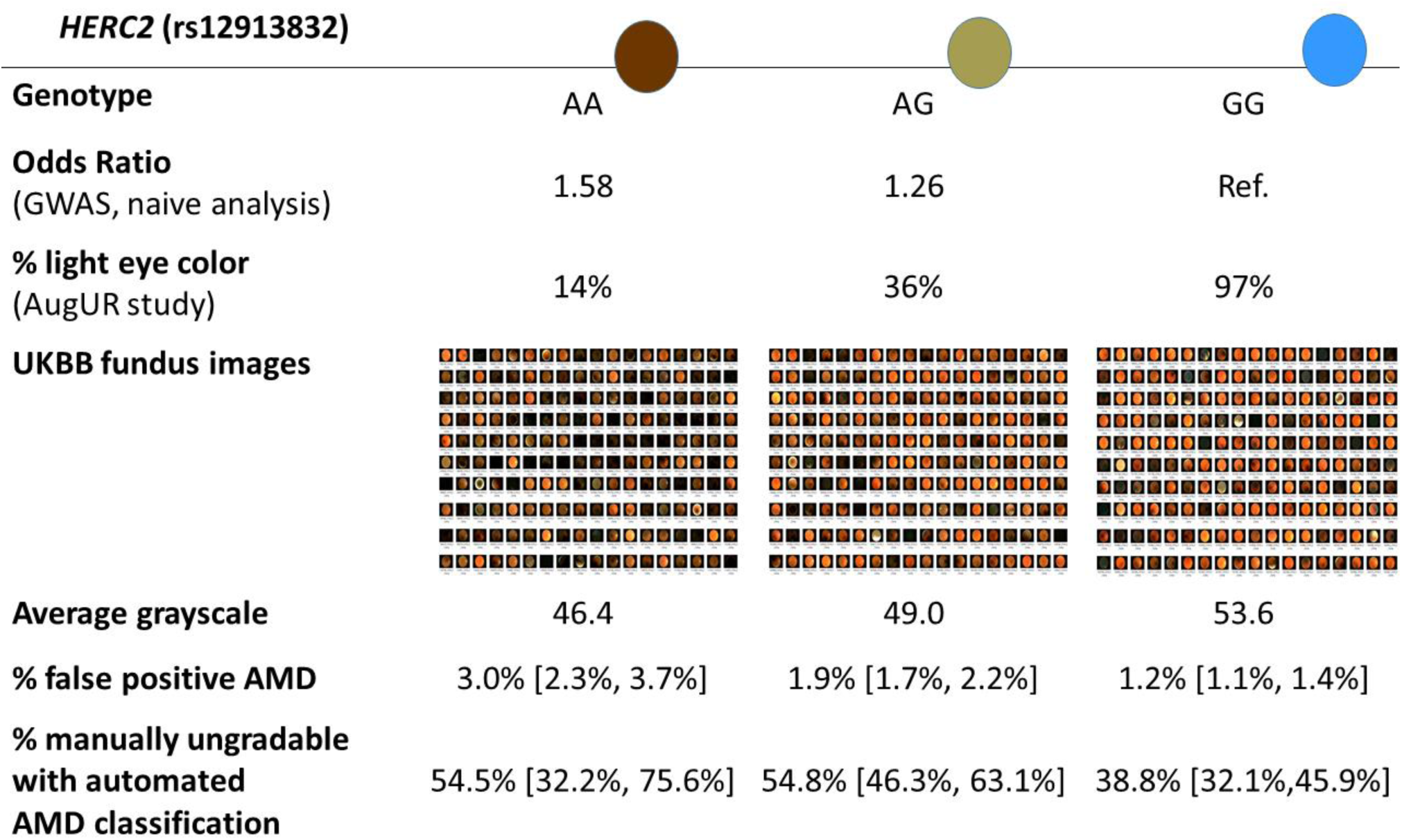
Evidence for differential misclassification in automatically derived AMD with respect to the *HERC2* variant rs12913832. Shown are (i) estimated odds ratios from the naïve analysis ignoring misclassification and various characteristics per genotype group: (ii) the fraction of persons with self-reported “light eye color” in the AugUR study, (iii) randomly selected fundus images in UKBB, (iv) image-lightness quantified by mean average grayscale, (v) proportion of false-positive AMD in the automated classification (1-specificity) and 95% confidence interval estimated via MLA2, and (vi) observed proportion of manually ungradable images that were deemed gradable by the algorithm and classified as “any AMD” or “AMD-free”.

## DISCUSSION

GWAS on machine learning derived classification of imaging-based diseases, like AMD, can be expected to accelerate knowledge gain and drug target development^30^, since it will enable substantially increased sample sizes and refined, homogeneous phenotyping. To this date, there was no GWAS reported using a machine learning derived classification for AMD or any other imaging-based disease – to our knowledge. We here present a GWAS on machine learning derived AMD in UK Biobank highlighting chances and challenges. By this GWAS on AMD combined with an evaluation of emerging genetic signals via our newly developed MLA, we were able to detect known AMD loci and to distinguish true loci from artefacts.

Such artefacts, i.e. false positives, can derive from a misclassification that is associated with a genetic variant. Our data and analyses provide a compelling example for such an artefact: our MLA revealed the *HERC2* signal as false positive signal and suggested darker eye color and darker fundus images as a relevant source of misclassification for this machine learning algorithm. It is perceivable that the misclassification process of other algorithms for AMD and for other image-based diseases will depend on one or the other characteristic as well, and that such a characteristic is picked up by some genetic variants due to the abundant range of genetically pinpointed characteristics (see e.g. NHGRI-EBI GWAS Catalog^31^), which can yield artefact signals when left unaccounted.

Our MLA, developed for bilateral diseases, does not only quantify the misclassification and the dependencies, but also guards against bias and artefacts in association analyses. Similar approaches are available for classic diseases^7,8^. Thus, this concept can be generalized to other algorithms and other image-based diseases. Our work here links the theory of misclassification to machine learning derived disease classification, which can be generalized also to measurement error and quantitative phenotypes.

We recommend a GWAS combined with a post-GWAS evaluation of emerging genetic effects for non-differential and differential misclassification not only to search for GWAS signals on image-based, machine-learning derived disease phenotypes. We also recommend such a GWAS as a quality control for diseases like AMD, where strong genetic signals are known: a GWAS on AMD ascertained by any classification approach, manual or automatic, should be able to detect at least the two strong known signals around *ARMS2/HTRA1* and *CFH*. When a GWAS does not detect these signals, this indicates issues that can be anything from mis-matched bio-samples, analytical errors, or imperfect disease ascertainment – like from machine learning algorithms as highlighted here. A GWAS can be a quick guide towards phenotype classification quality when genomic data is available.

Overall, we illustrate chances and challenges of machine learning derived disease classification in GWAS, and the applicability of our MLA to guard against bias and artefacts.

## Appendices

### Appendix A. MLA to adjust for response misclassification in bilateral diseases

We developed an MLA to adjust for response misclassification from an error-prone, entity-specific disease classification in bilateral diseases. Here we illustrate it based on the example of age-related macular degeneration, where AMD can occur in each eye (eye-specific AMD) and the person-specific binary outcome is defined as worse-eye outcome, i.e. “AMD in at least one eye”, and modeled using logistic regression. We assume that we have an error-prone, eye-specific AMD classification (e.g. from a machine-learning based automated classification) available for nearly all eyes and true, gold-standard classifications (e.g. manual classification) for a subset of individuals from validation data.

Let (Z_1i_, Z_2i_) ∈ {0,1} be the true, binary disease stages in the two eyes of study participant i, i.e. (Z_1i_ = 1, Z_2i_ = 0) means that participant i suffers from AMD in the left eye and is unaffected from AMD in the right. When estimating the association of person-specific risk factors with AMD, one often defines a binary person-specific disease status as worse-entity AMD,Y_i_ ≔ max(Z_1i_, Z_2i_), Z_1i_, Z_2i_ ∈ {0,1}, and uses logistic regression to estimate the association of some covariates X with AMD: the person-specific disease status Y_i_ equals 1, if at least one eye of individual i is classified as AMD, and Y_i_ equals 0, if both eyes are unaffected. As described previously^16^, such a worse-eye disease status can be misclassified because of two reasons: either, because of missing disease information in one of two eyes (in this case disease can be overlooked), or because of error-prone disease status for any of the two eyes. Here we assume that we observed an error-prone, eye-specific disease status 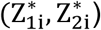 for each of the two eyes of a “main study” participant i and additionally the true disease status in each of the two eyes (Z_1j_, Z_2j_) for a subset of study participants j from the “validation study”. For all participants from the main study (error-prone classifications only) or the validation subset (error-prone and true classification), there is the additional issue that the disease information can be missing in one of two eyes, because of missing or ungradable fundus images. Since the automated (error-prone) and manual (gold standard, “true”) classification may judge differently on whether an image is gradable or ungradable, any possible subset of 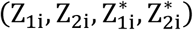 might be the available information for a specific study participant. To obtain valid estimates for the association of covariates with the true AMD status, we set up a likelihood based on the conditional probabilities of the observed error-prone and/or true eye-specific disease classifications given covariates. The product of these conditional probabilities over all individuals forms the likelihood, which has to be numerically optimized with respect to the regression parameters to obtain estimates. The different likelihood contributions for the individuals depend on the available AMD classifications (true and/or error-prone for one or both eyes).

The general problem of response misclassification when AMD information is missing in one of two eyes and/or the eye-specific classification suffers from misclassification with known classification probabilities has already been evaluated in a previous publication^16^. There, we also derived the corresponding likelihood contributions for the different scenarios of available outcome data. Here, we add the aspect that validation data is available for some study participants or, more specifically, a collection of error-free (gold-standard) classified single eyes, and that we model the eye-specific misclassification process based on information from this validation data. In the following, we describe the general idea and provide formulas for the respective likelihood contributions:

The assumed logistic regression model for the true worse-eye disease corresponds to the assumption that max(Z_1i_, Z_2i_) = Y_i_∼Bernoulli(π_i_), where we model the success probability based on a linear predictor via 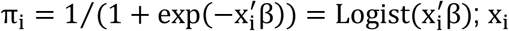 is a vector of observed person-specific covariates and β the vector of corresponding regression coefficients. It follows that P(Y_i_ = 1|x_i_) = π_i_. If we focus on single-eye disease classifications, there exist four different pattern of true disease classifications (Z_1i_, Z_2i_): (1,1), (1,0), (0,1), (0,0). From the assumed logistic regression model for Y_i_, it follows that P(Z_1i_ = 0, Z_2i_ = 0|x_i_) = 1 − π_i_. Based on the law of total probability, we can derive P(Z_1i_ = 1, Z_2i_ = 1|x_i_) = P(Z_1i_ = 1, Z_2i_ = 1|x_i_, Y_i_ = 1) × P(Y_i_ = 1|x_i_) and we define the person-specific conditional probability of being affected by AMD in both eyes given AMD in at least one eye as δ_i_ ≔ P(Z_1i_ = 1, Z_2i_ = 1|x_i_, Y_i_ = 1). When assuming symmetric probabilities for disease in one but not the other eye for left and right eyes (i.e. same probabilities to be affected in the left but not the right eye and vice versa), the conditional probability mass function of the two-entity disease status distribution can be written concisely as

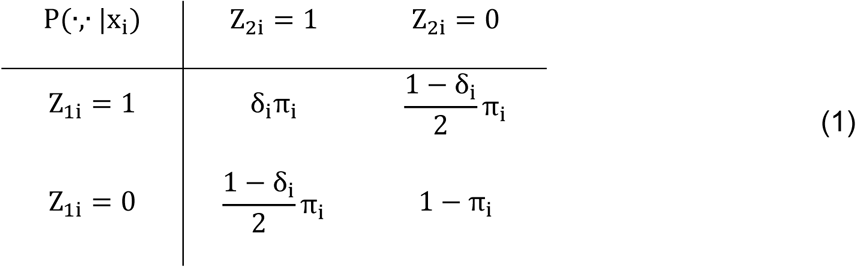

which specifies the *true data model*. If we look at a single eye selected randomly from both eyes, we can derive (without loss of generality for Z_1i_):

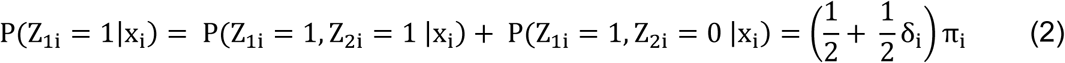

We now assume that we observed potentially misclassified single eye disease stages 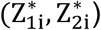 for each participant and describe the *misclassification process* based on the sensitivity and specificity of the classification,

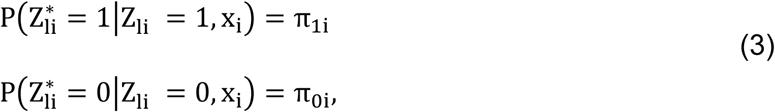

with l = 1,2; π_1i_ and π_0i_ are the person-specific sensitivity and specificity from the eye-specific classification process. We assume that the eye-specific classification process within an individual is independent in the two eyes, i.e.:

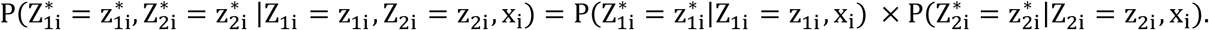

Based on the *true data model* and the description of the *misclassification process* via sensitivity and specificity, we can now express the conditional probabilities of all combinations of observed outcomes, by using Bayes’ rule and the law of total probability. If all four AMD classifications were observed for an individual (individual with full validation data, true and error-prone disease status for each of the two eyes), we can derive the following (omitting a random variable notation and only using the small z’s for the observed data):

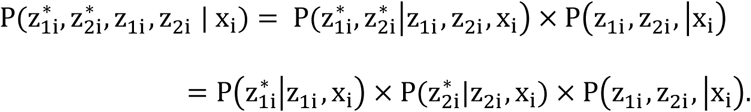

Here, we fraction the conditional probability of the observed data into terms of the eye-specific classification process (depending on sensitivity or specificity when the observed true outcome z_li_ is 1 or 0, respectively, (3)) and the true data model (1). If only the two eye-specific error-prone classifications are observed (individual in the main study, not part of the validation subset), the law of total probability can be used and the conditional probability can be expressed as

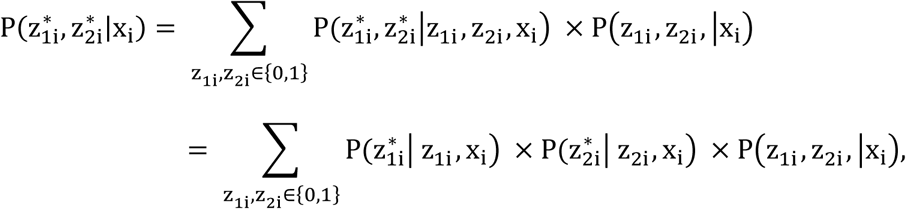

This again yields an expression that depends on the eye-specific classification probabilities (3) and the *true data model* (1).

If only a classification for one error-prone outcome was observed (e.g. 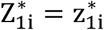), the conditional probability is given by

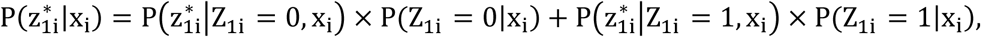

where the first terms in each summand depends on the specificity and the sensitivity of the eye-specific observation process; an expression for the second was already given above (equation (2)).

When three classifications were observed, e.g. 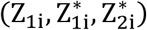 or 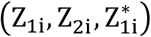, we can derive

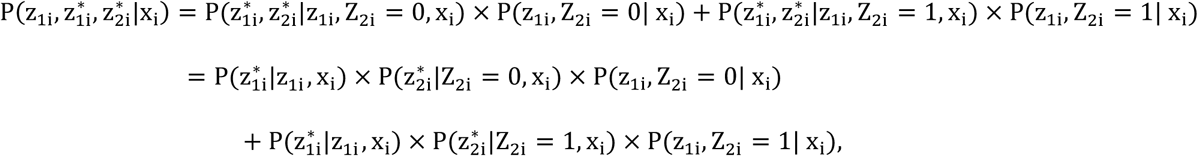

and

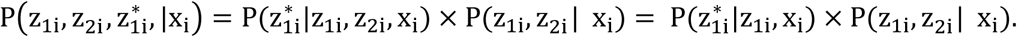

All conditional probabilities characterizing the *true data model* and the *misclassification process*, i.e. (i) the probability of true worse-eye AMD P(Y_i_ = 1|x_i_) = π_i_, (ii) the probability of AMD in both eyes given AMD in at least one eye P(Z_1i_ = 1, Z_2i_ = 1| Y_i_ = 1, x_i_) = δ_i_, (iii) the eye-specific sensitivity 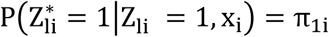 and (iv) the eye-specific specificity of the error-prone classification 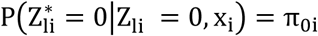, can potentially vary with person-specific characteristics. We therefore decided to model them based on the logistic function of a linear predictor, where relevant covariates (characteristics) can be specified for each probability. Combining all these expressions, we can set up the whole likelihood based on the derived conditional probabilities and numerically optimize with respect to the regression coefficients of the linear predictors for π_i_, δ_i_, π_1i_, and π_0i_. Standard errors of the maximum likelihood estimates are derived based on standard likelihood theory from the square root of the diagonal elements of the inverse of the observed Fisher information (Hessian) and used for inference. An implementation of the MLA in the statistical programming language R^32^ is available (**Web Resources**)

### Appendix B. Simulation study to evaluate consequences of ignoring misclassification and the performance of the MLA in correcting it

We performed a simulation study to evaluate the consequences of ignoring response misclassification and to evaluate the performance of the derived MLA in data scenarios similar to the situations in AMD studies. For each simulation scenario (data generating process), we simulated 1000 datasets, applied different models to the sampled data and evaluated the distribution of effect estimates, frequencies of significant statistical tests and coverage frequencies of confidence intervals for a central covariate of interest.

To sample data mimicking studies on AMD with internal validation data, we performed the following steps:

1. We sampled the true binary “worse-eye” AMD data Y for 5000 individuals by sampling from a Bernoulli distribution, where we modelled the success probability based on the logistic function of a linear predictor (corresponding to the assumed data generating process in logistic regression). For the linear predictor, we used an intercept of −0.25 (corresponding to an average probability of person-specific AMD of ∼0.44) and a continuous standard normal covariate X. We varied the log OR of X on Y between zero (simulation under H_0_ of no effect) and one.
2. To create the true eye-specific disease data (two binary observations per individual, (Z_1_, Z_2_)) we specified the conditional probability of being affected in both eyes given disease in at least one eye (i.e. Y = 1 based on “worse-eye definition), δ, to be (on average) δ = 1/(1 + exp(−1)) = 0.73. We assumed this probability to be either constant or varying with the continuous covariate X based on formula δ = 1/(1 + exp(−(1 + 1 × X))) = Logist(1 + 1 × X). For all individuals with sampled Y=1, we sampled a Bernoulli variable based on probability δ, to decide whether they were affected in both eyes or not. If they were affected on only one eye, we sampled randomly from the left or right.
3. To mimic the situation of missing information in one of two eyes, we sampled a Bernoulli random variable for each individual based on a fixed success probability (e.g. 0.75), to indicate whether information on both eyes was available. If not, we removed the disease information from a randomly selected eye.
4. To obtain eye-specific error-prone outcome data 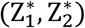, we conditioned on the true, sampled observations (Z_1_, Z_2_), and sampled the error-prone outcomes based on specified classification probabilities, the sensitivity P(Z* = 1|Z = 1) and specificity P(Z* = 0|Z = 0). Sensitivity and specificity were either fixed (non-differential misclassification, e.g. sens=spec=0.9) or varying between individuals based on the formula sens = Logist(2.20 + β_sens_ × X) for different values of β_sens_ (analogously for the specificity, corresponding to an average sens=spec=0.9).
5. Afterwards, we split the data into two parts, the “main study” and the “validation” subset based on defined fractions (e.g. n^val^ = 1000, n^main^ = 4000). For the validation subset we kept both, the true and the error-prone eye-specific AMD observations 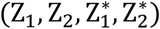; for the main study, we kept only the error-prone outcomes 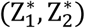 (or only the respective information for one of the two eyes, when information in one eye was missing for an individual).
6. For the naïve analysis ignoring response misclassification, we defined an observed, binary naïve person-specific outcome 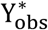 the following way: for individuals from the validation data, we used the true eye-specific disease information; for individuals from the main study data, we used the error-prone eye-specific information. When disease information was available for both eyes, we defined 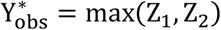 or 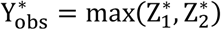, respectively; for observations with information only on one eye Z_1_, we used 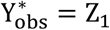 or 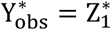. For individuals from the validation data with information on both eyes, 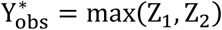 corresponds to the true Y; for all others, 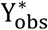 might be misclassified.

For each sampled dataset we estimated three models: 1) standard logistic regression based on the error-prone naïve worse-entity outcome 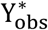, 2) the derived MLA (see above) modelling the probability of person-specific AMD and the probability of AMD in both eyes given AMD in at least one eye, δ, based on covariate X, while assuming a constant eye-specific sensitivity and specificity and accounting for missing information in one of two eyes (MLA1), and 3) the derived MLA allowing for a dependency of sensitivity and specificity on X (MLA2).

### Appendix C. Power analysis for reported lead variants based on UK Biobank sample size

We wanted to evaluate the impact of using the MLA on selected variants including the 34 reported lead variants known for their association with advanced AMD. Given reported effect sizes and effect allele frequencies (EAF), we expected the power to detect some of these 34 associations to be limited in a sample size of approximately 3,500 cases (and more controls). Therefore, we aimed to assess the power to detect reported genetic associations for AMD in the available data of UK Biobank, to focus our analyses with the MLA only on adequately powered reported associations and to avoid overinterpreting results from underpowered analyses. It is, however, not fully straight forward how to compute power for the scenario of “any AMD” from machine learning based disease classification, due to the power-diminishing effect of misclassification and some uncertainty of what effect size to use. We chose to use the reported^10^ EAFs in advanced AMD cases and AMD-free controls for the established 34 lead variants and computed the power for a t-Test on EAFs for differently sized groups, given the 3,544 cases and 44,521 controls derived from the automated “any AMD” classification in the UK Biobank GWAS data (**Supplemental Table 2**). The standard error of the difference in EAFs between cases and controls was derived based on the formula

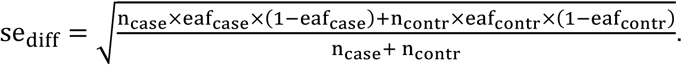

Based on these power calculations, we selected all lead variants with at least 80% power to yield nominally significant associations in UK Biobank. By this, we made the assumptions that EAFs in advanced AMD cases are transferable to EAFs of “any AMD” cases and that no misclassification was present in the machine learning derived any AMD classification. Therefore, this is probably an overestimate of available power. We performed the power analysis, however, mainly to dismiss variants with an obvious lack of power, while trying to include as many variants as reasonable in our analyses using the MLA.

### Appendix D. MLA avoids bias and excess of type-I error in simulation studies

In our simulation study, we investigated bias and type-I error of logistic-regression based association estimates for a binary worse-entity outcome Y ≔ max(*Z*_1_, *Z*_2_) ∈ {0,1} and a continuous covariate X, when error-prone single-entity observations 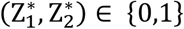 are observed instead of the true entity-specific disease classifications (Z_1_, Z_2_) ∈ {0,1}. When utilizing the error-prone observations for deriving the worse-entity outcomes 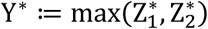, the entity-specific misclassification is passed on to the worse-entity disease stage. We compare the performance of the naïve analysis (logistic regression ignoring misclassification) and the two versions of our MLA for different simulation scenarios.

In the naïve analysis, we found a similar pattern for bilateral disease misclassification as reported for classic diseases^5,7^: (i) under the null hypothesis (**Table 1, Supplemental Table 1**, β_Y_ = 0), we found biased estimates and a lack of type-I error control (potential for false-positive association findings) for differential misclassification. With non-differential misclassification, estimates were unbiased and type-I error frequencies were at the desired levels. (ii) When X was associated with true AMD (**Table 1, Supplemental Table 1**, β_Y_ = 1), effect estimates were biased towards the null for non-differential misclassification and into any direction for differential misclassification. Specific for the bilateral disease situation was (iii) increasing bias with increasingly missing AMD in one of the two eyes, and (iv) a larger bias by decreased specificity than by decreased sensitivity. (**Table 1, Supplemental Table 1**).

In logistic regression, the larger the misclassification probabilities, the larger the bias of estimates^5^, with similar influence of increased probabilities for false-positive and false-negative classifications for balanced data. In the following, we provide an explanation of the findings (iii) and (iv) for bilateral diseases from above. Finding (iii) is explained by the fact that an increased fraction of missing eyes implies a reduced sensitivity for person-specific AMD: AMD in the missing eye can be overlooked, which can lead to a false-negative person-specific AMD classification if only the missing eye of an individual is affected. Finding (iv) was that decreased specificity had larger impact on bias than decreased sensitivity, e.g. for (sens, spec)=(0.9, 0.9) and a fraction of 25% of individuals with “missing eyes” and a true log OR of X on Y of 1 the observed bias was −0.27. When the sensitivity was reduced to 0.8 (specificity=0.9), the bias increased (in absolute value) to −0.32; when the specificity was reduced to 0.8 (sensitivity=0.9), the bias increased to −0.39. This can be explained by rewriting the probability of misclassification in the worse-entity outcome, P(Y* ≠ Y) as

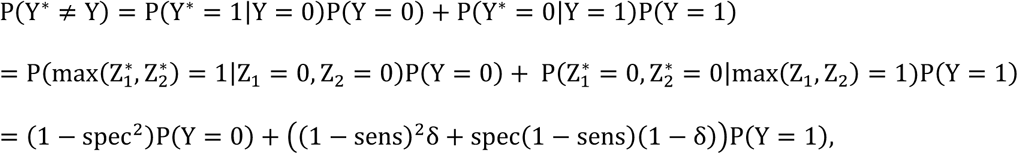

This illustrates the dependency of P(Y* ≠ Y) on entity-specific sensitivity, specificity, probability of disease in both entities given disease in one eye δ, and the fraction of truly affected individuals P(Y = 1). This probability can be evaluated for different combinations of parameters: for example, in the simulation study, we assumed P(Y = 1) = 0.44, δ = 0.75 (**Appendix B**), which leads to a misclassification probability of 12%, 14%, or 22% for (sens, spec)=(0.9, 0.9), (sens, spec)=(0.8, 0.9), or (sens, spec)=(0.9, 0.8), respectively, illustrating the larger impact of reducing specificity. This is even more true in scenarios with a lower fraction of affected individuals: if we assume a probability of person-specific disease of 0.10 instead of 0.44, we obtain misclassification probabilities of 17%, 18%, or 33%, for the same combinations of sensitivity and specificity. A reduced entity-specific specificity increases the probability of falsely classifying healthy entities towards disease, and falsely classifying only one of two healthy entities towards disease is sufficient to misclassify the person-specific disease status.

When applying the MLA1, we found it to effectively correct for bias and to yield the expected confidence interval coverage rates (∼95%) when the misclassification was non-differential, but we found it to still result in biased estimates and excess type-I error when the misclassification was differential (**Table 1, Supplemental Table 1**). When applying the MLA2, we found it effective in bias correction and type-I error control under all misclassification scenarios, but with larger standard errors due to the larger number of parameters in the model (**Table 1, Supplemental Table 1**). Overall, our simulation results documented substantial bias and lack of type-I error control when the naïve analysis was applied to misclassified data and our MLA to effectively remove bias and keep type-I error when specified correctly.

### Appendix E. Detailed results of MLA for the selected 26 variants

For estimating sensitivity and specificity, we found the following: (i) for the 3 lead variants from this GWAS (*CFH, ARMS2/HTRA1*, or *HERC2*, respectively), the MLA1-derived sensitivity and specificity (at mean age and two copies of the non-effect allele) showed only small differences between the 3 variants (sensitivity = 65%, 67%, 63%; specificity=98%, 98%, 99%, respectively, **Supplemental Table 6a**). From a model without including a genetic covariate, we obtained an overall sensitivity of 64.5% (95%-CI: 60.1%, 68.7%) and a specificity of 98.6% (98.4%, 98.8%). (ii) We did not find strong evidence for associations with age using MLA1 or MLA2 based on any of the 26 selected variants, except for an association of the specificity with age based on MLA1 for the *HERC2* variant that disappeared when applying MLA2 (age-P=6.71×10^−9^ or 0.70, respectively, **Supplemental Table 6a**). (iii) Applying MLA2, we found no association of the sensitivity with any selected variant (P>0.05/(23*2)), but a strong association of the specificity with the *HERC2* lead variant rs12913832 and with the reported *CFH* lead variant rs10922109 (OR_spec_=0.64, P_spec_=7.38×10^−9^ and OR_spec_=1.36, P_spec_=2.29×10^−4^, respectively; **Supplemental Table 6**).

Second, we obtained genetic association estimates from MLA1 and MLA2 accounting for misclassification and compared these with naïve analysis estimates. We found interesting patterns: (i) when applying MLA1, we found comparable, slightly increased effect estimates for the *CFH, ARMS2/HTRA1*, and *HERC2* lead variant when compared to the naïve analysis (MLA1: OR=1.23, 1.48, 1.34; P=1.69×10^−6^, 8.9×10^−18^, 1.11×10^−12^; naïve: OR=1.14, 1.30, 1.26, P=6.18×10^−7^, 2.44×10^−20^, 4.16×10^−16^; **Figure 2, Supplemental Table 7a**). (ii) When applying MLA2, we found similar effect estimates for *CFH and ARMS2*/*HTRA1* compared to MLA1 and naïve analysis (OR=1.19 or 1.28, respectively), which is in line with limited bias due to differential misclassification. We also found larger P-values (P=0.02 or 2.47×10^−4^, respectively, which is in line with larger uncertainty when estimating more model parameters. In contrast, we found a completely vanished effect estimate for the *HERC2* variant (MLA2: OR=1.03, P=0.76; **Figure 2, Supplemental Table 7a**), indicating a bias in the naïve analysis and MLA1 when ignoring a differential misclassification. (iii) Effect estimates for the 3 reported *CFH* variants increased when applying MLA2 compared to the naïve analysis. This was particularly interesting for the reported *CFH* lead variant rs10922109, where we now observed a nominally significant association into the reported direction (MLA2: OR=1.15, P=0.047; naïve: OR=1.00, P=0.98; **Supplemental Table 7c**). This is in line with the observed association of the specificity with this *CFH* variant. (iv) For the other 20 reported lead variants, we found many variants with increased effect estimates by MLA1 or MLA2 compared to the naïve analysis; effect estimates were mostly more comparable to reported effect sizes for advanced AMD^10^ (**Figure 3c**). For one variant, this MLA2 analysis yielded an effect into the opposite direction compared to the reported effect direction, which is the *C9* lead variant (OR=0.83, P=0.59). With an effect allele frequency of 1%, it is the rarest analyzed variant of the 26 selected variants and estimates from the reported association as well as for the MLA2 analysis have low precision (i.e. large standard errors).

## Supplemental Data

Supplemental Data include one figure and seven tables.

## Acknowledgements

This work was funded by DFG HE 3690/5-1 (to I.M.H.) and NIH R01 RES511967 (to I.M.H.), the University of Regensburg and the Ludwig-Maximilians-University München. The UK Biobank (accessed via application number 33999) was established by the Wellcome Trust medical charity, Medical Research Council, Department of Health, Scottish Government and the Northwest Regional Development Agency. It has also had funding from the Welsh Assembly Government, British Heart Foundation and Diabetes UK.

## Web Resources

An open source R implementation of the MLA to account for misclassification in bilateral disease in genetic association analyses is available at: https://www.genepi-regensburg.de/MLA-bilateral/ (upon publication)

Convolutional Neural Net Ensemble used for automated AMD classification and recommendations by the authors: https://github.com/RegensburgMedicalImageComputing/ARIANNA;

IrfanView: https://www.irfanview.com/;

GWAS catalogue: https://www.ebi.ac.uk/gwas/

## References

1. Litjens, G., Kooi, T., Bejnordi, B.E., Setio, A.A.A., Ciompi, F., Ghafoorian, M., van der Laak, J.A.W.M., van Ginneken, B., and Sánchez, C.I. (2017). A survey on deep learning in medical image analysis. Med. Image Anal. 42, 60–88.

2. Moreno-Torres, J.G., Raeder, T., Alaiz-Rodríguez, R., Chawla, N. V., and Herrera, F. (2012). A unifying view on dataset shift in classification. Pattern Recognit. 45, 521–530.

3. Csurka, G. (2017). A comprehensive survey on domain adaptation for visual applications. In Domain Adaptation in Computer Vision Applications, (Springer), pp. 1–35.

4. Heinze-Deml, C., and Meinshausen, N. (2017). Conditional variance penalties and domain shift robustness. ArXiv Prepr. 1710.11469.

5. Neuhaus, J. (1999). Bias and efficiency loss due to misclassified responses in binary regression. Biometrika 86, 843–855.

6. Hausman, J.A., Abrevaya, J., and Scott-Morton, F.M. (1998). Misclassification of the dependent variable in a discrete-response setting. J. Econom. 87, 239–269.

7. Carroll, R.J., Ruppert, D., Stefanski, L.A., and Crainiceanu, C.M. (2006). Measurement Error in Nonlinear Models (Chapman and Hall/CRC).

8. Lyles, R.H., Tang, L., Superak, H.M., King, C.C., Celentano, D.D., Lo, Y., and Sobel, J.D. (2011). Validation data-based adjustments for outcome misclassification in logistic regression: an illustration. Epidemiology 22, 589–597.

9. Klein, R., Meuer, S.M., Myers, C.E., Buitendijk, G.H.S., Rochtchina, E., Choudhury, F., de Jong, P.T.V.M., McKean-Cowdin, R., Iyengar, S.K., Gao, X., et al. (2014). Harmonizing the Classification of Age-related Macular Degeneration in the Three-Continent AMD Consortium. Ophthalmic Epidemiol. 21, 14–23.

10. Fritsche, L.G., Igl, W., Bailey, J.N.C., Grassmann, F., Sengupta, S., Bragg-Gresham, J.L., Burdon, K.P., Hebbring, S.J., Wen, C., Gorski, M., et al. (2016). A large genome-wide association study of age-related macular degeneration highlights contributions of rare and common variants. Nat. Genet. 48, 134–143.

11. Bycroft, C., Freeman, C., Petkova, D., Band, G., Elliott, L.T., Sharp, K., Motyer, A., Vukcevic, D., Delaneau, O., O’Connell, J., et al. (2018). The UK Biobank resource with deep phenotyping and genomic data. Nature 562, 203–209.

12. Burlina, P.M., Joshi, N., Pekala, M., Pacheco, K.D., Freund, D.E., and Bressler, N.M. (2017). Automated Grading of Age-Related Macular Degeneration From Color Fundus Images Using Deep Convolutional Neural Networks. JAMA Ophthalmol. 135, 1170.

13. Ting, D.S.W., Cheung, C.Y.-L., Lim, G., Tan, G.S.W., Quang, N.D., Gan, A., Hamzah, H., Garcia-Franco, R., San Yeo, I.Y., Lee, S.Y., et al. (2017). Development and Validation of a Deep Learning System for Diabetic Retinopathy and Related Eye Diseases Using Retinal Images From Multiethnic Populations With Diabetes. JAMA 318, 2211.

14. Grassmann, F., Mengelkamp, J., Brandl, C., Harsch, S., Zimmermann, M.E., Linkohr, B., Peters, A., Heid, I.M., Palm, C., and Weber, B.H.F. (2018). A Deep Learning Algorithm for Prediction of Age-Related Eye Disease Study Severity Scale for Age-Related Macular Degeneration from Color Fundus Photography. Ophthalmology 125, 1410–1420.

15. Peng, Y., Dharssi, S., Chen, Q., Keenan, T.D., Agrón, E., Wong, W.T., Chew, E.Y., and Lu, Z. (2019). DeepSeeNet: A Deep Learning Model for Automated Classification of Patient-based Age-related Macular Degeneration Severity from Color Fundus Photographs. Ophthalmology 126, 565–575.

16. Günther, F., Brandl, C., Heid, I.M., and Küchenhoff, H. (2019). Response misclassification in studies on bilateral diseases. Biom. J. 61, 1033–1048.

17. McCarthy, S., Das, S., Kretzschmar, W., Delaneau, O., Wood, A.R., Teumer, A., Kang, H.M., Fuchsberger, C., Danecek, P., Sharp, K., et al. (2016). A reference panel of 64,976 haplotypes for genotype imputation. Nat. Genet. 48, 1279–1283.

18. Walter, K., Min, J.L., Huang, J., Crooks, L., Memari, Y., McCarthy, S., Perry, J.R.B., Xu, C., Futema, M., Lawson, D., et al. (2015). The UK10K project identifies rare variants in health and disease. Nature 526, 82–89.

19. Keane, P.A., Grossi, C.M., Foster, P.J., Yang, Q., Reisman, C.A., Chan, K., Peto, T., Thomas, D., Patel, P.J., and UK Biobank Eye Vision Consortium (2016). Optical Coherence Tomography in the UK Biobank Study - Rapid Automated Analysis of Retinal Thickness for Large Population-Based Studies. PLoS One 11, e0164095.

20. Davis, M.D., Gangnon, R.E., Lee, L.-Y., Hubbard, L.D., Klein, B.E.K., Klein, R., Ferris, F.L., Bressler, S.B., Milton, R.C., and Age-Related Eye Disease Study Group (2005). The Age-Related Eye Disease Study severity scale for age-related macular degeneration: AREDS Report No. 17. Arch. Ophthalmol. (Chicago, Ill. 1960) 123, 1484–1498.

21. Loh, P.-R., Kichaev, G., Gazal, S., Schoech, A.P., and Price, A.L. (2018). Mixed-model association for biobank-scale datasets. Nat. Genet. 50, 906–908.

22. Kutalik, Z., Johnson, T., Bochud, M., Mooser, V., Vollenweider, P., Waeber, G., Waterworth, D., Beckmann, J.S., and Bergmann, S. (2011). Methods for testing association between uncertain genotypes and quantitative traits. Biostatistics 12, 1–17.

23. Devlin, A.B., Roeder, K., and Devlin, B. (2013). Genomic Control for Association. 55, 997–1004.

24. Machiela, M.J., and Chanock, S.J. (2015). LDlink: a web-based application for exploring population-specific haplotype structure and linking correlated alleles of possible functional variants. Bioinformatics 31, 3555–3557.

25. Ma, C., Blackwell, T., Boehnke, M., Scott, L.J., and GoT2D investigators (2013). Recommended joint and meta-analysis strategies for case-control association testing of single low-count variants. Genet. Epidemiol. 37, 539–550.

26. Sturm, R.A., Duffy, D.L., Zhao, Z.Z., Leite, F.P.N., Stark, M.S., Hayward, N.K., Martin, N.G., and Montgomery, G.W. (2008). A single SNP in an evolutionary conserved region within intron 86 of the HERC2 gene determines human blue-brown eye color. Am. J. Hum. Genet. 82, 424–431.

27. Stark, K., Olden, M., Brandl, C., Dietl, A., Zimmermann, M.E., Schelter, S.C., Loss, J., Leitzmann, M.F., Böger, C.A., Luchner, A., et al. (2015). The German AugUR study: study protocol of a prospective study to investigate chronic diseases in the elderly. BMC Geriatr. 15, 130.

28. Brandl, C., Zimmermann, M.E., Günther, F., Barth, T., Olden, M., Schelter, S.C., Kronenberg, F., Loss, J., Küchenhoff, H., Helbig, H., et al. (2018). On the impact of different approaches to classify age-related macular degeneration: Results from the German AugUR study. Sci. Rep. 8, 8675.

29. Chakravarthy, U., Wong, T.Y., Fletcher, A., Piault, E., Evans, C., Zlateva, G., Buggage, R., Pleil, A., and Mitchell, P. (2010). Clinical risk factors for age-related macular degeneration: a systematic review and meta-analysis. BMC Ophthalmol. 10, 31.

30. Nelson, M.R., Tipney, H., Painter, J.L., Shen, J., Nicoletti, P., Shen, Y., Floratos, A., Sham, P.C., Li, M.J., Wang, J., et al. (2015). The support of human genetic evidence for approved drug indications. Nat. Genet. 47, 856–860.

31. Buniello, A., Macarthur, J.A.L., Cerezo, M., Harris, L.W., Hayhurst, J., Malangone, C., McMahon, A., Morales, J., Mountjoy, E., Sollis, E., et al. (2019). The NHGRI-EBI GWAS Catalog of published genome-wide association studies, targeted arrays and summary statistics 2019. Nucleic Acids Res. 47, D1005–D1012.

32. R Core Team (2019). R: A Language and Environment for Statistical Computing.

