## Supplemental Figure 1 for "Chances and challenges of machine learning based disease classification in genetic association studies illustrated on age-related macular degeneration"

**Supplementary Figures: Machine learning phenotypes in genetic association analysis as misclassification problem exemplified on age-related macular degeneration**

**Supplementary Figure 1. Estimated genetic associations for automatically derived “any AMD” in the *HERC2* region based on three models in UK Biobank participants.** Shown are region plots of genetic association P-values for 6,567 genetic variants based on three models: **a)** logistic regression adjusted for age and 2 genetic principal components (naïve analysis ignoring misclassification). **b)** as a) but accounting for a misclassification that depended on the *HERC2* lead variant (rs12913832, chr15:28365618) via MLA2. **c)** as b) but modelling the misclassification as depending on each respective *HERC2* region variant via MLA2.

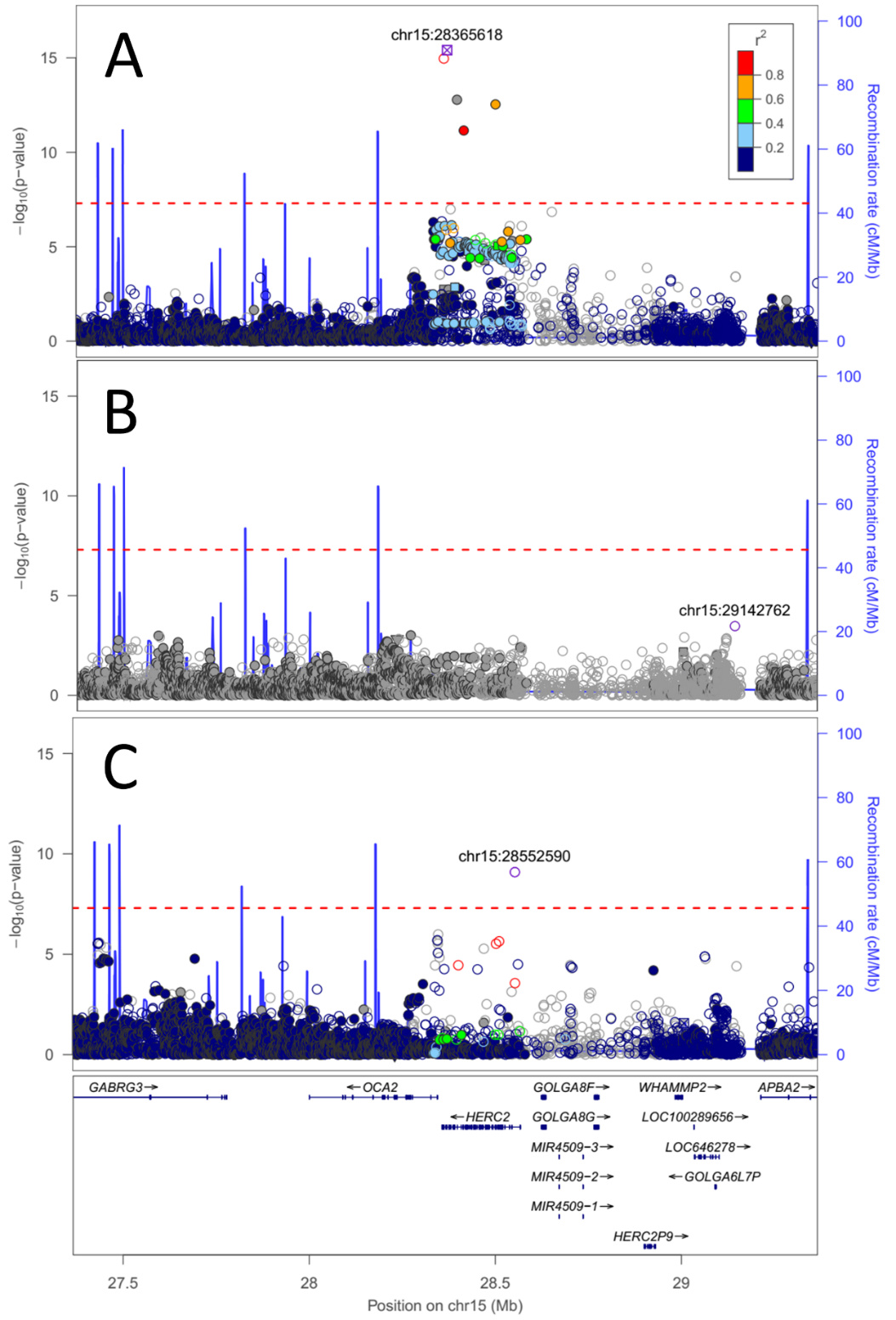
